# Reference-informed prediction of alternative splicing and splicing-altering mutations from sequences

**DOI:** 10.1101/2024.03.22.586363

**Authors:** Chencheng Xu, Suying Bao, Hao Chen, Tao Jiang, Chaolin Zhang

## Abstract

Alternative splicing plays a crucial role in protein diversity and gene expression regulation in higher eukaryotes and mutations causing dysregulated splicing underlie a range of genetic diseases. Computational prediction of alternative splicing from genomic sequences not only provides insight into gene-regulatory mechanisms but also helps identify disease-causing mutations and drug targets. However, the current methods for the quantitative prediction of splice site usage still have limited accuracy. Here, we present DeltaSplice, a deep neural network model optimized to learn the impact of mutations on quantitative changes in alternative splicing from the comparative analysis of homologous genes. The model architecture enables DeltaSplice to perform “reference-informed prediction” by incorporating the known splice site usage of a reference gene sequence to improve its prediction on splicing-altering mutations. We benchmarked DeltaSplice and several other state-of-the-art methods on various prediction tasks, including evolutionary sequence divergence on lineage-specific splicing and splicing-altering mutations in human populations and neurodevelopmental disorders, and demonstrated that DeltaSplice outperformed consistently. DeltaSplice predicted ∼15% of splicing quantitative trait loci (sQTLs) in the human brain as causal splicing-altering variants. It also predicted splicing-altering *de novo* mutations outside the splice sites in a subset of patients affected by autism and other neurodevelopmental disorders, including 19 genes with recurrent splicing-altering mutations. Among the new candidate disease risk genes, *MFN1* is involved in mitochondria fusion, which is frequently disrupted in autism patients. Our work expanded the capacity of *in silico* splicing models with potential applications in genetic diagnosis and the development of splicing-based precision medicine.

## INTRODUCTION

Most human genes consist of multiple exons and introns that are transcribed into precursor mRNAs (pre-mRNAs). The process of pre-mRNA splicing that removes introns and joins exons is required to produce mature mRNAs ready for protein translation (Black 2003; Chen and Manley 2009). In addition, alternative splicing allows single genes to generate multiple mRNA transcript isoforms, thus dramatically amplifying the complexity of genetic information encoded in the genome (Nilsen and Graveley 2010). Given the importance of accurate splicing for gene expression, it is not surprising that aberrant splicing has been implicated in an expanding list of genetic diseases, ranging from neurologic disorders to cancer (Cooper et al. 2009; Scotti and Swanson 2016). Determining splicing-altering mutations is thus an important arena to improving disease diagnosis. Substantial efforts have also been made to target splicing for precision medicine, such as correcting aberrant splicing and skipping exons that contain disease-causing mutations. In addition, manipulating alternative splicing, such as the suppression of endogenous “poison exons” containing premature termination codons (PTCs) and activation of cryptic splice sites can provide a powerful means of modulating gene expression using therapeutically compatible approaches (El Marabti and Abdel-Wahab 2021).

A longstanding question in the splicing field is to derive a “splicing code”, which can accurately predict splice sites and their quantitative usage from pre-mRNA sequences, including those containing genetic mutations (Black 2000; Wang and Burge 2008; Bao et al. 2019). This task has been challenging because accurate splicing is dictated by not only the universal splicing signals including 5 ▪ and 3 ▪ splice sites, a polypyrimidine tract, and a branch site, but also by numerous additional splicing-regulatory elements embedded in the exon and flanking intronic sequences (Black 2003; Chen and Manley 2009). These *cis*-regulatory signals, which individually contain limited information due to their size and degeneracy, must be recognized and integrated by hundreds of RNA-binding splicing factors to recruit or antagonize the splicing machinery. The abundance and activity of splicing factors vary in different cellular contexts, resulting in cell-type- or cell-state-specific splicing (Licatalosi and Darnell 2010; Ule and Blencowe 2019). While disruption of the invariant splice sites inevitably abolishes splicing, most mutations affecting splicing-regulatory elements change splice site usage (SSU) quantitatively. Therefore, accurate splicing prediction requires a computational algorithm to properly identify and weigh all the contributing splicing-regulatory sequences, which involve enormous combinatorial complexity.

Most earlier efforts to predict splicing used a list of manually curated features related to splicing, such as various splicing factor motif sites identified in previous analyses (Wang et al. 2004; Barash et al. 2010; Leung et al. 2014; Xiong et al. 2015; Jha et al. 2017; Zhang et al. 2018). These studies provided important global insights and unequivocally demonstrated that SSU is determined by both splice site strengths and the enrichment of different types of splicing-regulatory elements. However, the accuracy of these algorithms is often limited when applied to predict the usage of individual splice sites or exons. Since the last decades, deep sequencing technologies have made it possible to accurately determine splice sites and their usage on a genome-wide scale. Furthermore, deep learning methods provide a powerful engine to develop predictive models directly from genomic sequences without requiring manual feature engineering. In particular, applications of convolutional neural networks (CNNs) have led to remarkable progress in splicing prediction (Bretschneider et al. 2018; Zuallaert et al. 2018; Jaganathan et al. 2019; Louadi et al. 2019; Zeng and Li 2022; Celaj et al. 2023). Among them, SpliceAI uses a residual network architecture with dilated convolution to predict splice sites based on 10kb flanking sequences (Jaganathan et al. 2019). It achieved high prediction accuracy when tested with canonical transcripts and successfully predicted mutations causing cryptic splicing, including private mutations in human populations and *de novo* mutations affecting patients with autism and other neurodevelopmental disorders. However, this method was not designed to predict quantitative SSU explicitly, a limitation that was addressed by Pangolin (Zeng and Li 2022). Pangolin uses a similar network architecture as SpliceAI, but with several notable modifications. Instead of using binary labels of splice sites in canonical transcripts, where alternatively spliced weak splice sites are under-represented, Pangolin uses quantitative usage of all splice sites detected in the RNA-seq data of four different species to train the model. It also models SSU in multiple tissues simultaneously using multi-head output layers. These modifications were found to improve the prediction of splicing-altering mutations in human populations and diseases.

Despite these progresses, the current methods still exhibit limited accuracy in predicting SSU. In this study, we describe DeltaSplice, a deep learning model designed to enhance the *in silico* prediction of SSU and splicing-altering mutations. Compared to previous methods using deep learning models, DeltaSplice uses SSU data from eight mammalian species to train larger and more accurate models. Importantly, it incorporates paired gene sequences to fully exploit the similarity inherent in homologous sequences and learn the effects of mutations on splicing. The network architecture of DeltaSplice also enables the utilization of a reference sequence together with the corresponding SSU, which is frequently available in many practical applications, to improve the prediction accuracy on a target sequence (e.g., the same gene containing disease-associated mutations or a homologous gene in another species). We demonstrate that DeltaSplice consistently outperformed the other benchmarked state-of-the-art methods, including SpliceAI and Pangolin, in various prediction tasks.

## RESULTS

### DeltaSplice model overview

The DeltaSplice model consists of a feature extraction module and multiple prediction modules. It uses a residual convolutional neural net as its core building blocks, which is similar to SpliceAI and Pangolin. However, DeltaSplice has several important differences in network design to best predict the splice-site probability and SSU from pre-mRNA sequences. DeltaSplice uses deeper networks with larger receptive fields than previous work (30kb vs 10kb) to integrate more distal signals in longer genes (with 11.6 fold more model parameters than SpliceAI). A key feature of DeltaSplice is that it operates in two modes: the single-sequence mode and the dual-sequence mode (Fig. 1). In the single-sequence mode, the model is trained to predict the splice-site probability **p_s_** and SSU **u_s_** within individual input gene sequences, denoted as **s**. In the dual-sequence mode, DeltaSplice predicts the SSU **u_t_** for a target gene sequence, referred to as **s_t_**, by utilizing information from a reference gene sequence, denoted as **s_r_**, along with the corresponding reference SSU, denoted as **u_r_**. While the dual-sequence-mode model shares the model parameters of the feature-extraction module with the single-sequence mode, it has its own prediction modules. The dual-sequence mode is specifically designed to predict splicing-altering mutations when the SSU of the reference gene sequence is known. For example, the reference sequence can be the gene sequence of healthy control individuals, while the target sequence is the gene sequence containing mutations from patients affected by a genetic disease. The incorporation of the reference SSU together with the reference gene sequence enables DeltaSplice to improve the accuracy in predicting the impact of mutations when the SSU cannot be reliably predicted from the sequences alone.

**Figure 1:**
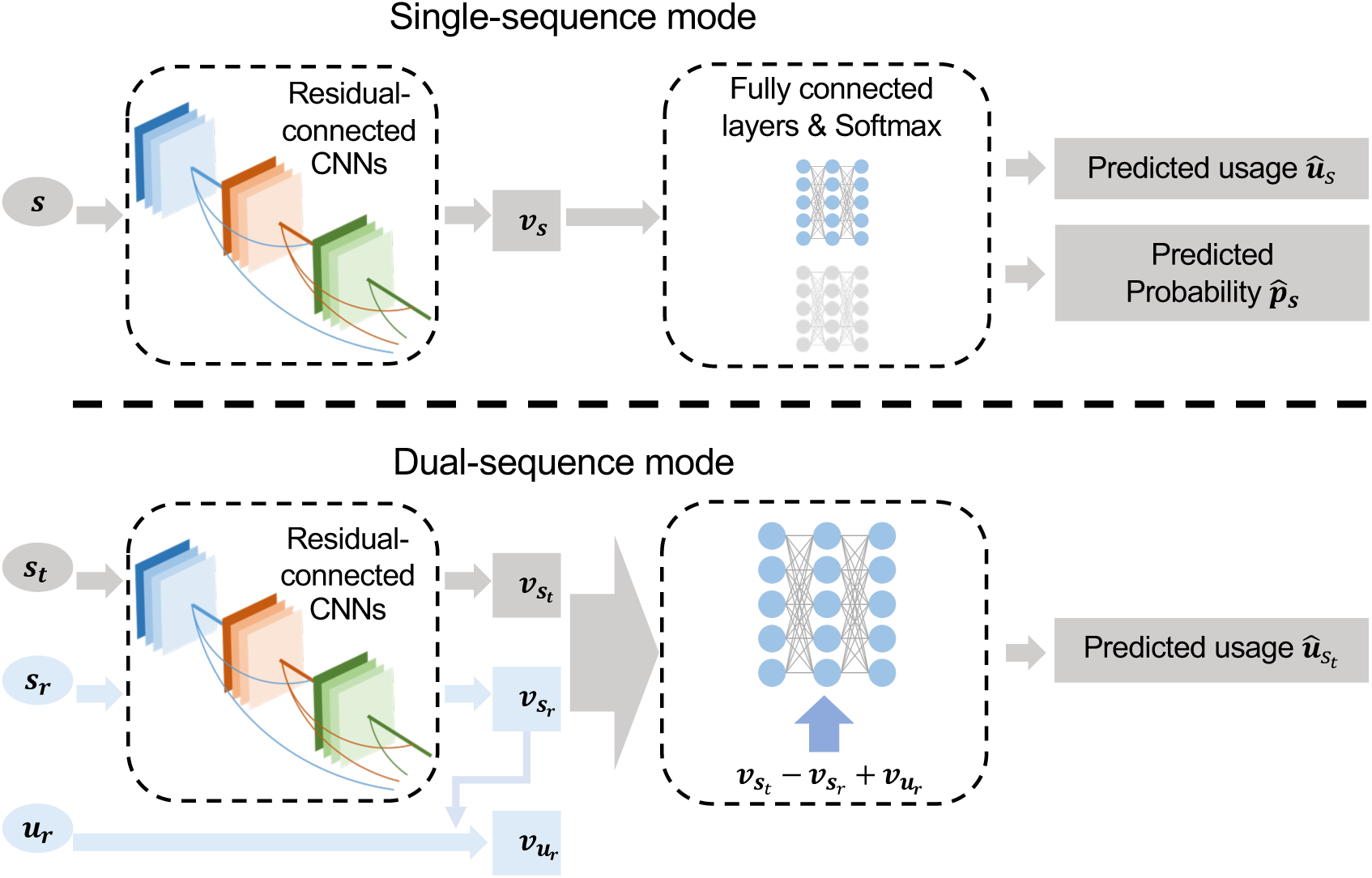
The architecture of DeltaSplice. DeltaSplice comprises a feature-extraction module and multiple prediction modules. The feature-extraction module is constructed using residual-connected convolutional neural networks (CNNs), which convert a one-hot encoded input sequence to a feature representation. Each prediction module consists of fully connected layers and a Softmax output layer that module takes a feature representation as input and generates predictions for SSU or splice site probabilities. The single-sequence mode employs two prediction modules to predict the SSU 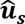 and the splice site probabilities 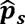 for each site in the input gene sequence s, based on the corresponding feature representation v_s_. In the dual-sequence mode, the feature-extraction module calculates the feature representation ***v*_*s*_t__** and ***v*_*s*_r__** separately for the target gene sequence ***s*_t_** and the reference gene sequence ***s*_r_**. The predicted SSU 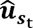 for every site in the target gene sequence is computed using a prediction module, from the input ***v*_*s*_t__ - *v*_*s*_f__ + *v*_*u*_r__**. Here ***v*_*u*_r__** is the feature representation of the reference SSU ***u*_r_**.

To train DeltaSplice models, we used splice sites and SSU in adult brain tissues quantified using RNA-seq data from eight different mammalian species, including human, chimpanzee, rhesus macaque, marmoset, cow, pig, mouse, and rat (Supplemental Tables S1 and S2). In the single-sequence mode, the training adopts a supervision approach similar to SpliceAI and Pangolin. The ideal training data for the dual-sequence mode are splicing-altering genetic variants or mutations, but unfortunately, no such large-scale mutation datasets with experimentally determined impacts are available. To overcome this limitation, DeltaSplice randomly samples pairs of gene sequences with varying degrees of similarities and their corresponding SSU to train the model and learn the impact of genetic mutations. Finally, a unified loss function is established by combining the supervision of both single-sequence and dual-sequence modes, enabling the model parameters to be trained for both modes simultaneously (for further details, see Methods).

### Quantitative prediction of SSU

We first benchmarked the performance of DeltaSplice in predicting SSU in the brain across various species using the single-sequence mode, in comparison with SpliceAI and Pangolin. To assess how data from multiple species influence the performance of DeltaSplice, we also trained an intermediate model using only human genes and their SSU, which was denoted as DeltaSplice (human). The distributions of SSU values predicted by DeltaSplice (trained with data from eight species) and DeltaSplice (human) more closely align with the experimental SSU distributions, showing higher correlations than those predicted by SpliceAI and Pangolin (Fig. 2A,B). This improvement suggests the benefits of longer input sequences and larger models. Moreover, DeltaSplice model trained with all species data achieved further improved performance as compared to the model trained with human data alone, suggesting the benefit of the expanded dataset. This observation was also made recently with Pangolin (Zeng and Li 2022).

**Figure 2:**
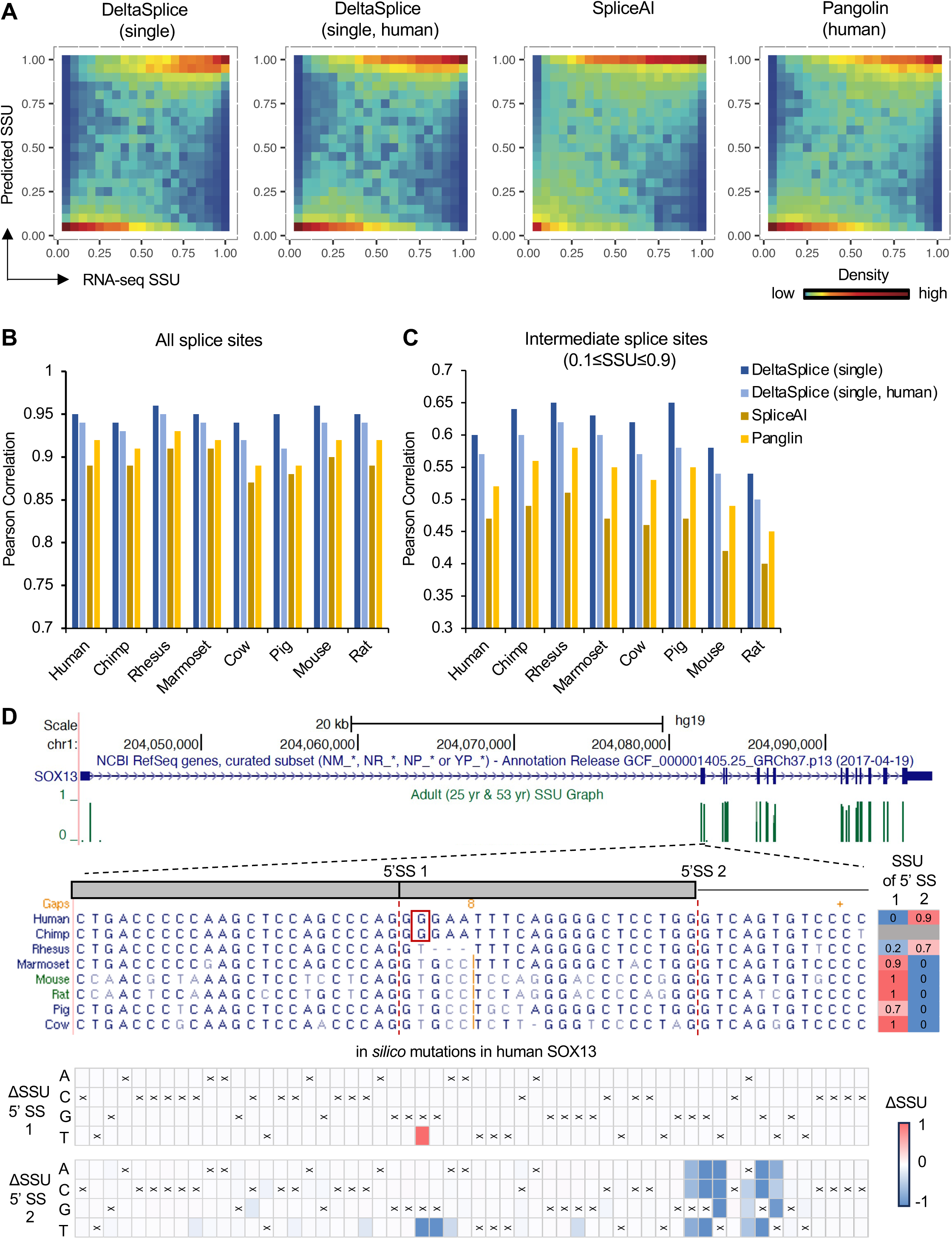
The performance of DeltaSplice in single-sequence mode and baseline methods in predicting SSU across different species. **A**. Distribution of human SSU predicted by different methods in comparison with experimental measurements by RNA-seq. DeltaSplice (single) refers to DeltaSplice in single-sequence mode trained with data from all species, while DeltaSplice (single, human) was model in single-sequence mode trained using only human data. SpliceAI and Pangolin were trained with human data. **B**. Pearson correlation between RNA-seq and predicted SSU for all splice sites. **C**. Similar to (B) but for splice sites with experimental SSU between 0.1 and 0.9. **D**. DeltaSplice predicts mutations driving lineage-specific splicing in *SOX13*. *SOX13* exon 2 has two alternative 5 ▪ splice sites, as shown in the gene schematics. Multiple-sequence alignments of the alternatively spliced region are shown, together with the usage of the two splice sites in each species on the right. Note that there is insufficient RNA-seq read coverage to quantify SSU in chimpanzees, although the limited number of reads supports the use of 5 ▪ splice site 2. The impact of *in silico* mutations in each nucleotide position in the region on the two splice sites is shown in the heatmaps at the bottom. The nucleotide base in each position of the human gene sequence is indicated by ‘x’.

SSU is bimodally distributed (Yan et al. 2015), with only ∼20% of splice sites exhibiting SSU within the 0.1 to 0.9 range in our dataset. The accuracy of SSU prediction for these intermediate splice sites is crucial in delineating the impact of splicing-altering mutations outside the splice sites, which led us to focus our evaluation specifically on these splice sites. For all compared methods, the prediction of weak and strong splice sites (SSU<0.1 or >0.9, respectively) is in general quite accurate, while quantitative prediction of SSU of intermediate splice sites is more challenging (Fig. 2A-C and Supplemental Fig. 1). Nevertheless, DeltaSplice has a distinct advantage over SpliceAI and Pangolin in predicting SSU of intermediate splice sites, archiving a correlation of 0.6 in human, as compared to 0.47 for SpliceAI and 0.52 for Pangolin (Fig. 2C and Supplemental Fig. 1). We also noticed that the splice-site probability scores predicted by SpliceAI tend to overestimate SSU, especially for weak and intermediate splice sites (SSU<0.5), as compared to DeltaSplice and Pangolin (Supplemental Fig. S1), suggesting the advantage of training models using quantitative data than binary labels.

We delved into a particularly interesting example predicted by DeltaSplice, focusing on *SOX13*, a member of SOX (SRY-related HMG-box) family transcription factors involved in embryonic development and cell lineage specification (Schepers et al. 2002). SOX13 protein is an autoantigen in type 1 diabetes, islet cell antigen 12 (ICA12) (Kasimiotis et al. 2001). We found that *SOX13* exon 2 (the first coding exon) has two alternative 5 ▪ splice sites showing lineage-specific splicing. Human, chimpanzee and rhesus macaque predominantly or exclusively use the downstream 5 ▪ splice site 2, resulting in a seven amino acid extension in an intrinsically disordered region of the encoded protein as compared to the other species analyzed in this study, which exclusively use the upstream 5 ▪ splice site 1 (Fig. 2D, upper panel). The 5 ▪ splice site 1 is disrupted in human and chimpanzee due to a T>G mutation (i.e., splice site dinucleotide GT>GG), which is consistent with the lineage-specific switch to the use of splice site 2. However, it remains unclear whether this substitution is the only driver mutation leading to the splicing divergence. DeltaSplice correctly predicted the usage of 5 ▪ splice site 2 in human (SSU=0.95 as compared to 0.93 as measured in RNA-seq; note SSU<1 due to partial intron retention). *In silico* mutations confirmed that mutations in 5 ▪ splice site 2 motif resulted in abrogation of its usage, as one would expect. Importantly, a G>T mutation that restores 5 ▪ splice site 1 in human *SOX13 in silico* was also predicted to abrogate the use of splice site 2 (SSU<0.01), while activating the usage of splice site 1 (SSU=0.97) (Fig. 2D, bottom panel). These results support the notion that this single nucleotide change likely played a critical role in the lineage-specific splicing and the introduction of the seven amino acids extension in the protein in humans and other closely related primates.

### Prediction of lineage-specific splicing changes

Given the imperfect accuracy of all compared methods, including DeltaSplice in single-sequence mode, in the quantitative prediction of SSU, we reasoned that the dual-sequence mode utilizing the SSU of the reference gene sequence, i.e., reference-informed prediction, may improve the prediction of splicing changes caused by mutations in human populations or related species during evolution. To test this hypothesis, we first evaluated the dual-sequence mode of DeltaSplice using homologous splice sites in the test set between humans and the seven other species. This evaluation aimed to assess DeltaSplice’s ability to predict changes in SSU (ΔSSU) between humans and each of the other compared species. In this analysis, gene sequences from humans are used as reference sequences, and the corresponding SSU is used as reference usage. For simplicity, the model in dual-sequence mode is referred to as DeltaSplice hereafter, while the model in the single-sequence mode included for comparison is denoted as DeltaSplice (single). Consistent with the higher accuracy of DeltaSplice (single) in predicting SSU of the intermediate splice sites, ΔSSU predicted by DeltaSplice in both dual- and single-sequence modes exhibit higher correlations with RNA-seq measurements, as compared to SpliceAI and Pangolin (Fig. 3A,B and Supplemental Fig. 2). Furthermore, in comparison of the dual- vs. single-sequence mode of DeltaSplice, the dual-sequence mode achieved a higher top *k* accuracy for predicting splicing differences between chimpanzee and human or between rhesus macaque and human. In this metric, each model was asked to predict the top *k* candidates where *k* is the true number of orthologous splice site pairs with validated splicing changes (|ΔSSU|≥0.2 as measured by RNA-seq) (Fig. 3C). However, the advantage of the dual-sequence mode was less clear when the human gene reference was used to predict splicing in more distal species from our analysis.

**Figure 3:**
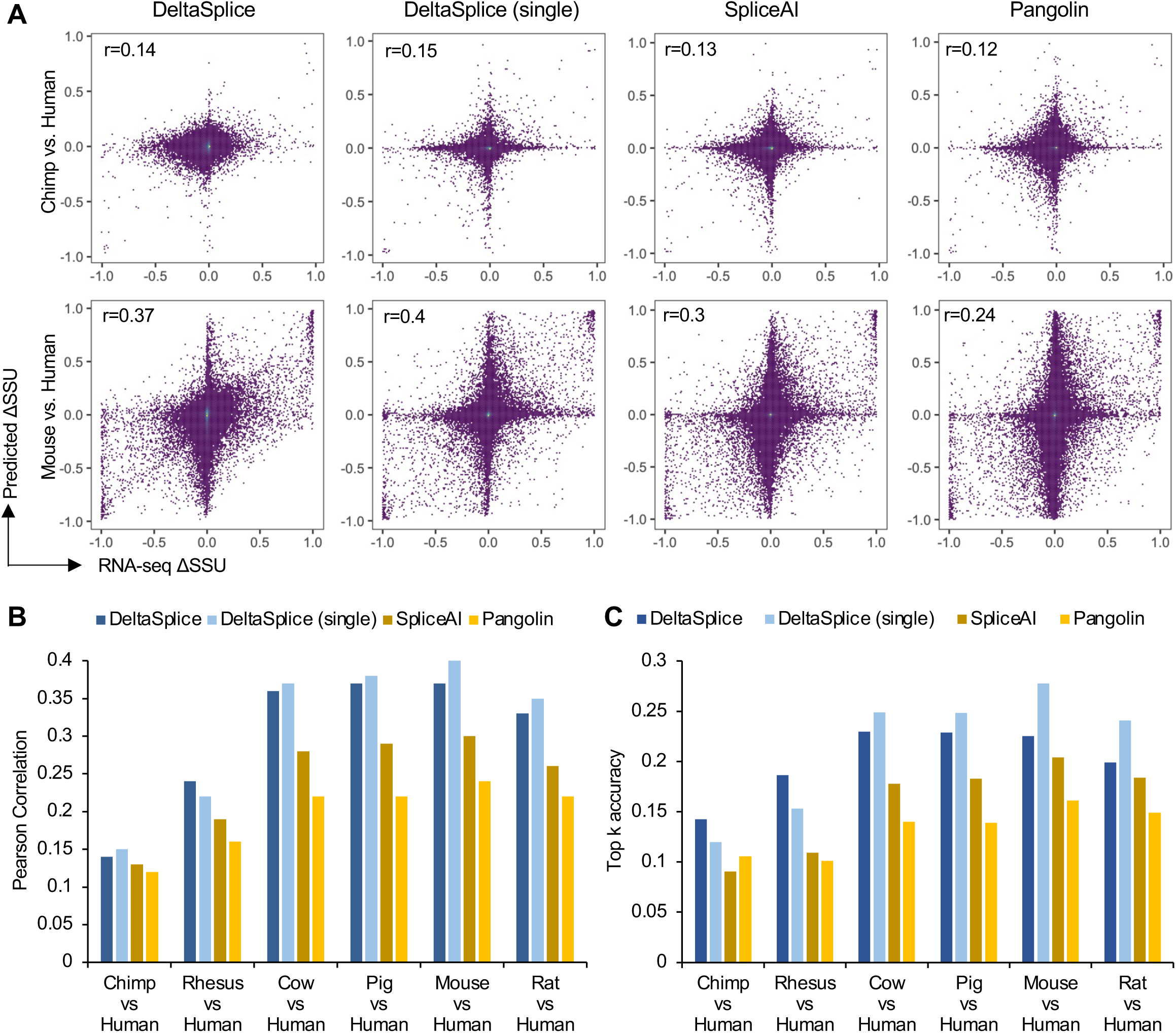
The performance of DeltaSplice and baseline methods in predicting ΔSSU of splice sites homologous to humans. **A**. Distribution of ΔSSU between chimpanzee and human (top), and between mouse and human (bottom). In each panel, ΔSSU measured by RNA-seq is shown on the x-axis and ΔSSU predicted by different methods is shown on the y-axis. DeltaSplice refers to the results obtained using the dual-sequence mode while DeltaSplice (single) refers to the results obtained using the single-sequence mode. **B**. Pearson correlation between RNA-seq and predicted ΔSSU of splice sites homologous to humans using different methods. **C**. Top *k* accuracy of different methods in predicting splice sites with |ΔSSU|≥0.2, where *k* is the true number of exons showing the specified differences.

### Prediction of splicing-altering mutations quantified by splicing reporter assays

To further investigate whether reference-informed predictions can improve the discovery of splicing-altering mutations, we next tested DeltaSplice and the other compared methods using multiple datasets derived from high-throughput splicing reporter assays. These datasets include the impact of single nucleotide variants (SNVs) on cassette exon inclusion as measured by Vex-seq (Adamson et al. 2018) and MFAS (Cheung et al. 2019), as well as the combinatorial effect of multiple SNVs on the inclusion of *FAS* exon 6 (Baeza-Centurion et al. 2019). In these prediction tasks, changes in percent exon inclusion (ΔPSI) were calculated by averaging ΔSSU of the two alternative splice sites and compared with corresponding experimental measurement. Another neural network model named MMSplice, which was particularly designed to predict splicing-altering mutations (Cheng et al. 2019), was included for comparison.

For the Vex-seq dataset, predictions by DeltaSplice and MMSplice showed a higher correlation with RNA-seq measurement than SpliceAI and Pangolin (r=0.69-0.71 vs. 0.56 for SpliceAI and Pangolin; Fig. 4A). For the *FAS* exon 6 dataset, DeltaSplice has a similar performance as SpliceAI, while predictions by Pangolin and MMSplice showed a somewhat lower correlation (Supplemental Fig. S3). The MFASS data provide a less quantitative measure of splicing changes derived from a fluorescent reporter-based readout. Therefore, the performance of each model was evaluated using a top *k* accuracy in predicting splicing-disrupting variants (SDVs) defined by ΔPSI<-0.5 between alternative and reference alleles. Again, DeltaSplice outperformed all other compared methods, with a top *k* accuracy of 0.53 as compared to 0.46-0.49 (where *k*=1,048 is the number of true SDVs in the dataset) (Fig. 4B). Notably, the advantage of DeltaSplice over the other compared methods was most evident when the analysis focused solely on SNVs more distal from the splice sites (i.e., > 20bp), with a top *k* accuracy of 0.39 vs. 0.25-0.31 (Fig. 4C).

**Figure 4:**
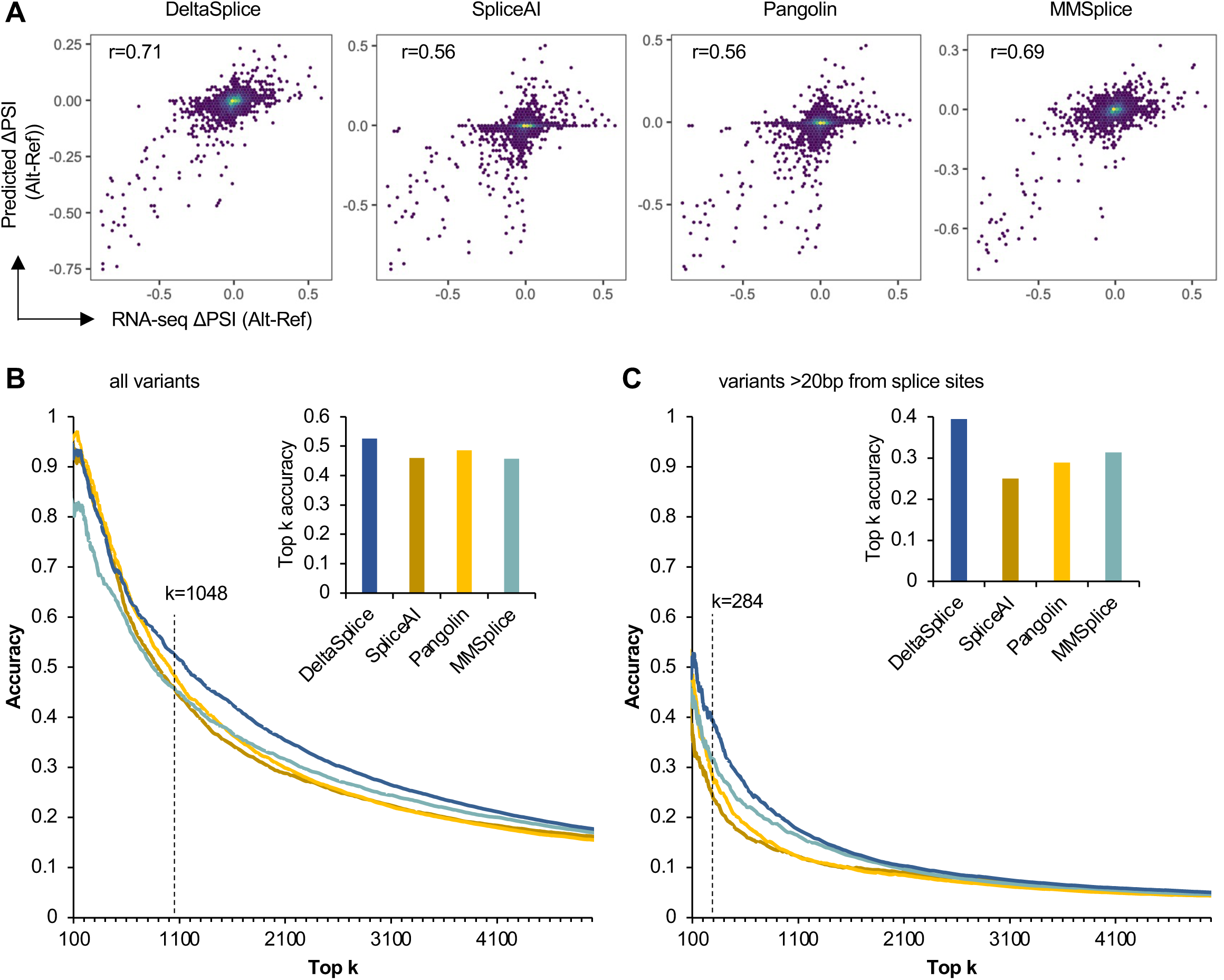
The performance of DeltaSplice and baseline methods in predicting splicing-altering mutations as measured by reporter assays. **A.** The distribution of ΔPSI measured by RNA-seq and predicted by each method is shown for SNVs in Vex-seq dataset. **B**. Top *k* accuracy of DeltaSplice and baseline methods on MFASS dataset. Splicing-disrupting variants (SDVs, which have ΔPSI_alt-ref_<-0.5) were used to calculate top *k* accuracy with varying *k*. The inset shows the top *k* accuracy with *k*=1048 (the true number of SDVs in the dataset). **C**. Similar to (B) but only for variants ≥20bp from the nearest splicing sites.

### Prediction of causative variants with splicing QTLs

Previous studies, such as those conducted by the Gene-Tissue Expression GTEx consortium, have identified numerous SNVs associated with splicing variation in human populations (splicing quantitative trait loci or sQTLs) (GTEx Consortium 2020). However, whether these sQTLs represent causal splicing-altering variants is unknown. We predicted the impact of brain sQTLs and control SNVs on splicing using DeltaSplice and the other compared methods by reasoning that splicing-altering SNVs should be enriched in sQTLs, and the intersections of the two methods can help to identify causal splicing-altering mutations given the complementarity of the two approaches. Indeed, top candidates predicted by different models all showed varying degrees of enrichment, and the largest enrichment was observed in predictions by DeltaSplice (4.1 fold as compared to 3.4-3.6 fold among top 20,000 predictions) (Fig. 5); excluding splice site mutations did not notably change the results (Supplemental Fig. 4). In total, DeltaSplice identified 891 brain sQTLs (representing 154 unique SNVs) with predicted |ΔPSI|>0.05, accounting for about 15% of all brain sQTLs (Supplemental Table S3).

**Figure 5:**
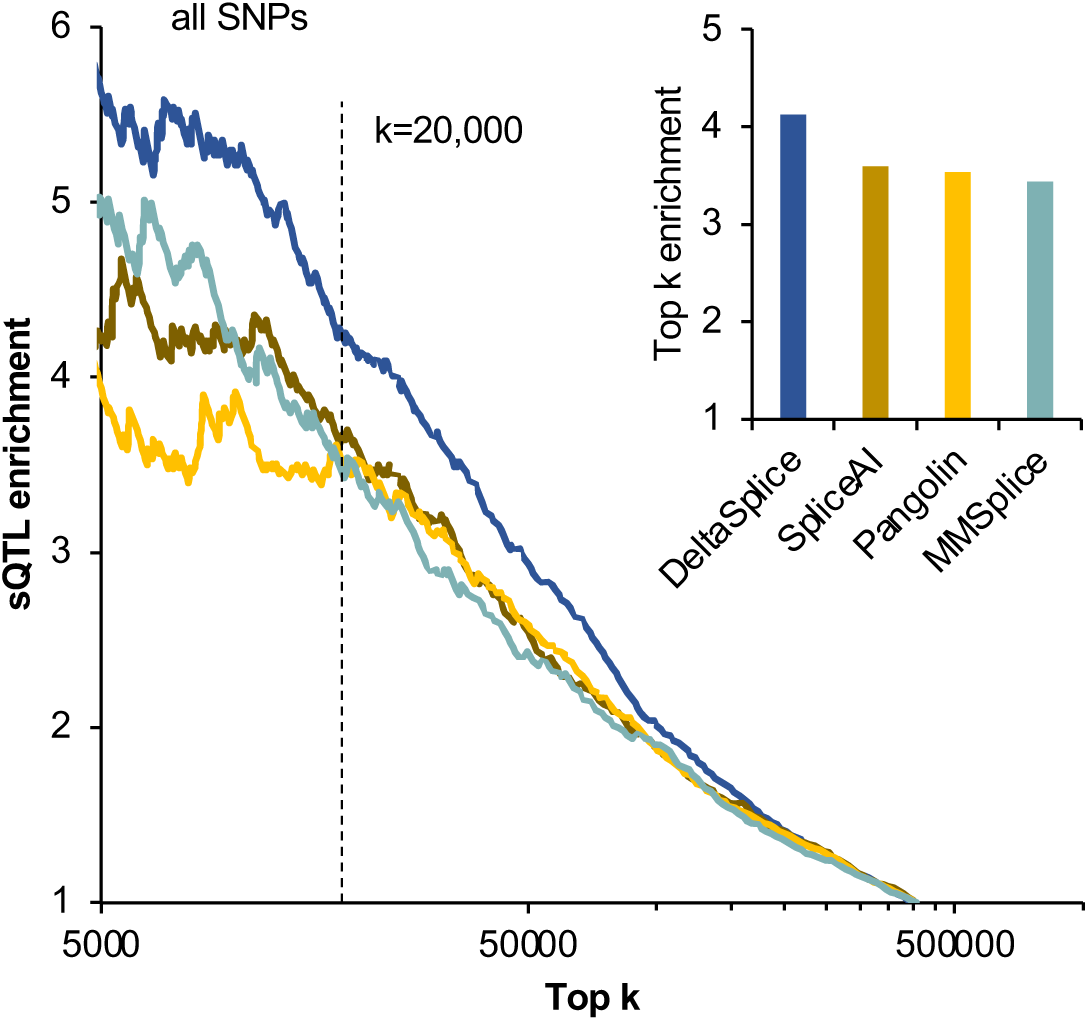
The performance of DeltaSplice and baseline methods in predicting splicing-altering mutations at sQTLs. GTEx brain sQTLs were used as foreground while common SNPs were used as background. Only variants ≤200bp from cassette exons were used for analysis. Foreground and background variants were combined and ranked by |ΔPSI| predicted by each method and the enrichment of sQTLs among top *k* predictions with varying *k* is shown. The inset shows the enrichment of sQTLs among top 20,000 variants.

### Prediction of *de novo* mutations implicated in autism and neurodevelopmental disorders

Gene-disrupting mutations, including splicing-disrupting mutations, by *de novo* mutations (DNMs) are important pathoetiologic mechanisms underlying neurodevelopmental disorders (Ronemus et al. 2014; Jaganathan et al. 2019; Dawes et al. 2023). However, previous analysis of splicing-disrupting mutations is largely limited to mutations in the known or cryptic splice sites, while those outside splice sites remain difficult to identify. We compared DeltaSplice and other methods in predicting splicing-altering mutations focusing on DNMs that are not stop gain/loss mutations and outside the splice sites. Note that this is more stringent than the previous analysis which did not exclude splice-site mutations (Jaganathan et al. 2019). For this analysis, we compiled a comprehensive list of DNMs identified by whole genome sequencing (WGS) and whole exome sequencing (WES) (Supplemental Tables S4 and S5). The WGS dataset consists of cases of autism spectrum disorders (ASDs) and unaffected controls. In this dataset, we observed that splicing-altering mutations predicted by DeltaSplice are enriched in autism patients (up to 1.4 fold; p=0.036, single-sided binomial test; Fig. 6A). Splicing-altering mutations predicted by other methods showed varying degrees of enrichment, but the magnitude is more moderate (Fig. 6A). Similar observations were made for DNMs associated with various neurodevelopmental disorders detected by WES (Fig. 6B). These results confirm that DeltaSplice has improved accuracy in predicting *bona fide* splicing-altering mutations outside the splice sites conferring disease risks.

**Figure 6:**
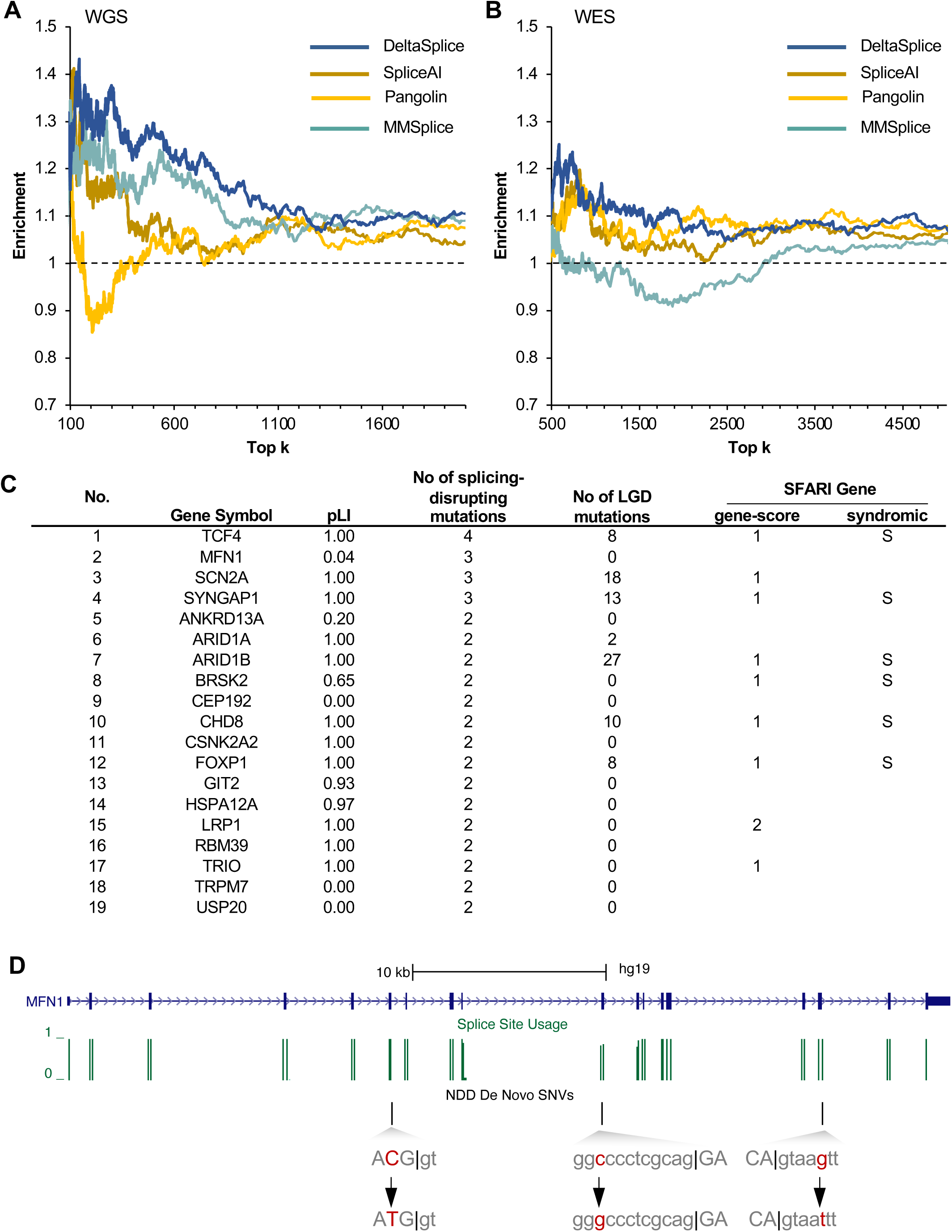
The performance of DeltaSplice and baseline methods in predicting splicing-altering *de novo* mutations associated with neurodevelopmental disorders (NDDs). **A**. Enrichment of autism-associated DNMs in the WGS dataset. Stop-gain/loss or splice-site mutations were excluded from this analysis. DNMs in autism patients or controls were combined and ranked by ΔPSI_alt-ref_. Among top *k* predictions using varying *k*, the enrichment of autism-associated mutations is shown. **B**. Similar to (A), but for DNMs identified in NDD and control samples in the WES dataset. **C.** Genes with ≥2 splicing-disrupting mutations (as defined by ΔPSI_alt-ref_ <-0.05). The number of likely gene-disrupting mutations (stop-gain or splice site mutations, but not frameshifting indels) as well as SFARI gene scores is also shown. **D**. Three predicted splicing-altering mutations in the *MFN1* gene. Exonic and intronic sequences around the mutations are shown in capital and small letters, respectively.

In total, we predicted 130 and 414 splicing-altering mutations (ΔPSI<-0.05 in alternative vs. reference alleles) in the WGS and WES datasets, accounting for 3.6% and 3.0% of patients respectively (Supplemental Tables S6 and S7). Intriguingly, after combining the prediction results from the two datasets and removing redundancy, we obtained 19 genes with two or more independent splicing-altering mutations (Fig. 6C and Supplemental Table S8). Among them, 8 genes have a confidence score of 1 (high-confidence) and one gene has a score of 2 (strong candidate) concerning their association with autism in the SFARI autism gene database (Abrahams et al. 2013). While a subset of genes in the list also have recurrent gene-disrupting mutations of other types (stop gain, frame-shifting insertions/deletions and splice site mutations), supporting their role in conferring autism risks, others do not have previous evidence from established mutation analysis. One interesting example is *MFN1*, which has three splicing-altering mutations predicted by DeltaSplice, with two mutations in ASDs (Satterstrom et al. 2020) and one mutation in intellectual disability (Lelieveld et al. 2016) (Fig. 6D). *MFN1* encodes a mitochondria membrane protein involved in mitochondria fusion, which has been reported to be disrupted in ASD patients. Interestingly, reduced expression of *MFN1* and several other genes involved in mitochondria fusion has been found in postmortem ASD brains, but the genetic basis of such alterations is unclear (Tang et al. 2013). Each of the three mutations in *MFN1* was predicted to reduce the inclusion of a constitutively spliced exon, which will result in a frameshift and thus reduction of the protein-coding mRNA transcripts. Our discovery of splicing-altering mutations suggests a possible genetic mechanism underlying certain mitochondria deficits in autism and other neurodevelopmental disorders.

## DISCUSSION

The ability to accurately predict splicing from primary sequences is not only a testimony of our understanding of the splicing code, but also has a wide range of practical applications. Tremendous progress has been made in the last few years by developing deep learning-based models and utilities of these models to facilitate clinical genetic diagnosis and development of precision medicine are on the horizon (Jaganathan et al. 2019; Zeng and Li 2022; Celaj et al. 2023). From our analysis, however, it seems to be clear that quantitative prediction of splicing changes, especially those involving alternative splice sites with intermediate strengths, remains a challenge. This study represents an effort to improve the current state of the art with several intuitive strategies. Given the enormous complexity of splicing regulation, we used an expanded dataset to allow training of larger models (11.6 fold more parameters as compared to SpliceAI) and integrating signals from longer genomic sequences. Since the number of human genes and splice sites is intrinsically limited, in contrast to images available for training of computer vision algorithms (Deng et al. 2009), a strategy we employed is to use data from multiple related species. This strategy was independently adopted by Pangolin (Zeng and Li 2022). We argue that the benefit of including homologous gene sequences for model training is more than an increase in sample size. Since homologous genes are located in the vicinities of human gene sequences in the high dimensional space, they are especially helpful in learning the impact of splicing-altering mutations, which also represent small perturbations in the human gene sequences. In our study, we designed a dual-sequence mode to optimize the learning from related genes. Another strategy that distinguishes our work from previous studies is to acknowledge that the current accuracy of quantitative SSU prediction is still less than ideal, so the SSU of the reference gene sequence, which is frequently readily available, can provide the correct baseline to improve the prediction of the altered SSU caused by mutations. We confirm the benefits of these strategies, as implemented in DeltaSplice, by benchmarking with the current cutting-edge methods in a range of prediction tasks. In particular, while SpliceAI and Pangolin were previously shown to predict cryptic splicing caused by mutations effectively, DeltaSplice demonstrated improved prediction of quantitative changes in SSU, which is more challenging than prediction of cryptic splice sites. This allowed us to identify additional splicing-altering mutations underlying splicing QTLs and associated with neurodevelopmental disorders, including autism.

We note that the current models do not lack room for improvement, and expect that further improvement can be made by optimizing the training data and network architecture. In addition, the current models do not explicitly model the cellular context such as the expression and activity of splicing factors. In contrast to the current models which can be only used to predict splicing in the same cellular context as the training data, a context-aware model would remove this restriction, thus enabling many perturbation experiments, such as splicing factor overexpression or depletion, to be performed *in situ* and expanding the use case of these computational splicing models.

## ACKNOWLEDGEMENT

We thank Zhang and Jiang lab members for helpful discussion throughout the project. This study was supported by grants from the National Institutes of Health (NIH) (R35GM145279 and P50HD109879 to CZ, and R01NS125018 to CZ and TJ). S.B. was in part supported by a Columbia Precision Medicine Research fellowship. High-performance computation was supported by NIH grants S10OD012351 and S10OD021764.

## AUTHOR CONTRIBUTIONS

Conceptualization and experimental design: CX, SB, TJ, CZ. DeltaSplice model development and implementation: CX, SB. Data analysis: CX, SB, HC, CZ. Writing: CX, SB, CZ. All authors critically reviewed the manuscript.

## METHOD

### Estimation of splice site usage in brain tissues across eight mammalian species

RNA-seq data from adult brain tissues of humans and seven other mammalian species were obtained from previous studies (see summary in Supplemental Table S1). The raw reads in fastq files were downloaded, mapped and processed to estimate splice site usage (SSU) in annotated genes. Specifically, RNA-seq reads for each sample were mapped using STAR (Dobin et al. 2013) to the reference genome including a list of annotated exon junctions. The following parameters were used for mapping: --alignSJoverhangMin 8 --alignSJDBoverhangMin 5 --outFilterMismatchNmax 999 --outFilterMismatchNoverReadLmax 0.06 --outFilterIntronMotifs RemoveNoncanonicalUnannotated. These parameters required a minimum of 5-nt overlap for known exon junctions and 8-nt overlap for novel exon junctions. Non-canonical exon junctions were allowed only if the exon junctions were previously annotated in the respective genes.

SSU is defined as the ratio of the exon junction reads spliced at the precise position over the total number of reads that cover the splice site (read counts from replicate samples were pooled). SSU is independent of constitutive or alternative splicing patterns. To estimate SSU, we used a comprehensive list of splice sites in annotated gene transcript models (such as RefSeq, UCSC Known Gene transcripts, Ensembl genes; see Supplemental Table S1), those previously detected in mRNA/EST sequences, and those identified in RNA-seq data (Yan et al. 2015). SSU was calculated for all 5□ or 3□ splice sites with a read coverage≥20 and estimated standard deviation ≤0.1, while sites with an insufficient number of reads were assigned ‘NA’ so that they would not be treated as non-splice sites during model training and testing.

Several practical issues were considered in the estimation of SSU. We decided to include multi-map reads in the calculation of SSU, as we found the exclusion of these reads appeared to result in more artifacts. We also examined the estimated SSU across all splice sites in each gene, and eliminated a gene unless the maximum SSU within the gene is ≥0.99. This filtering step was to exclude cases in which a gene is embedded in the intron of another gene, or there are spurious exon junction reads that span the whole gene, which can lead to the underestimation of SSU of all splice sites in the gene, including those constitutive splice sites with an expected SSU of 1.

To provide input for DeltaSplice, each input sequence of length *L* base pairs (bp) is one-hot encoded as a *L*×4 vector, while the SSU values are encoded by a *L*×3 vector, in which the three numbers of each nucleotide position represent the SSU as 3 ▪ splice site, 5 ▪ splice site, or non-splice site. To distinguish splice sites with low usage and non-splice sites, we set the minimum SSU of a *bona fide* splice site to be 1e-9. For DeltaSplice model training and testing, we used genes on human chromosomes 1, 3, 5, 7, and 9 to form a test dataset. We randomly selected one-third of the genes on the remaining chromosomes as a validation dataset to select model parameters, and the rest as the training dataset. For the other seven species, genes were assigned to training, validation, and test datasets based on the assignments of their orthologous genes in humans. This approach ensures that orthologous genes were assigned to the same dataset, to avoid information leaks between different datasets. The total number of genes and splice sites were summarized in Supplemental Table S2.

Each gene sequence is cut into sub-sequences with a sliding window of 5,000 bp. Each resulting sub-sequence, along with the 15,000 bp sequences both upstream and downstream of the sub-sequence (i.e., W=35,000 bp in total), serves as input for training the DeltaSplice model. DeltaSplice generates output predictions for the central 5,000 bp positions, while features were extracted from the flanking 30,000 bp around each position. In the evaluation process, DeltaSplice takes sequences of *n* bp (*n*>30,000) and provides predictions for the central *n*-30,000 bp.

### DeltaSplice network architecture

The architecture of DeltaSplice consists of a feature-extraction module and multiple prediction modules. The feature-extraction module is composed of residual-connected CNNs with 24 residual units, and each residual unit consists of two layers of dilated CNNs with 64 channels (i.e., 48 convolution layers × 64 channels). For comparison, SpliceAI (Jaganathan et al. 2019) uses 32 convolution layers × 32 channels. The one-hot encoded sequence is input to this module and transformed into a feature representation. Batch normalization (Ioffe and Szegedy 2015) is used to stabilize the model training and Dropout (Srivastava et al. 2014) is used to mitigate the overfitting issue in each residual unit. Each prediction module consists of three fully connected layers with ReLU activation functions and a Softmax layer. These modules calculate the predicted SSU or probability from the feature representation output by the feature extraction module. In total, DeltaSplice has 8,075,145 model parameters, which is 11.6 fold more than the number of parameters (698,787) in SpliceAI.

DeltaSplice operates in two modes: the single-sequence mode and the dual-sequence mode (Fig. 1). In the single-sequence mode, the feature-extraction module calculates the feature representation **v_s_** of the input sequence **s**. Subsequently, the model employs two prediction modules to independently predict SSU 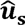 and the probability of being splice sites 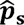 for each site in **s**. Both 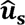 and 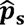 are a vector of three dimensions, representing the usage (probability) of the site as 3□ splice site, 5□ splice site, or non-splice site. In the dual-sequence mode, the feature-extraction module computes the feature representations **v_s_t__** and **v_s_f__** separately for the target sequence **s_t_** and the reference sequence **s_r_**. The reference SSU **u**_r_ is converted into its feature representation **v_u_r__** using a two-layer fully connected network with ReLU as the activation function. The model is constrained to learn feature representations in the feature space such that the difference between the feature vector of the input sequence and the feature representation of the corresponding SSU is minimized. The predicted SSU 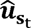 for every site in the target sequence is then calculated using a prediction module from **v_s_t__ - v_s_f__ + v_u_r__**.

### DeltaSplice model training

To effectively train DeltaSplice, we have devised four specific loss functions to optimize learning from the comparative analysis of gene pairs. To simplify the notation, we denote the cross-entropy between two matrices ***a*** and ***b***, both in the shape of W × 3, as

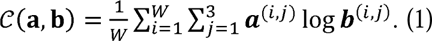

#### Classification loss ℓ_S,C_ in the single-sequence mode

The classification loss *ℓ*_S,C_ in the single-sequence mode is used to supervise DeltaSplice’s ability to accurately identify splice sites in gene sequences. The single-sequence mode predicts the SSU 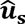 and the splice-site probability 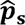 of each site in an input sequence **s**. The ground truth of 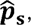 denoted by ***p*_s_**, is a binary matrix, in which each bit indicates if a position of **s** is a 3 ▪ splice site, 5 ▪ splice site, or neither. Then, *ℓ*_S,C_ is defined as the cross-entropy between ***p*_s_** and 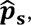

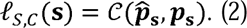

#### Regression loss ℓ_S,R_ in the single-sequence mode

*ℓ*_S,R_ is used to supervise DeltaSplice’s ability to accurately predict SSU in the single-sequence mode. Let ***u*_s_** denote the ground truth of 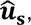 *i.e.*, the RNA-seq measured SSU of each site in the input gene sequence **s**. It is important to note that to incorporate splice-site types into ***u*_s_**, we let ***u*_s_** have the same shape as ***p*_s_**. Specifically, 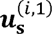 and 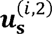 denotes the SSU of site if this site is a 3 ▪ or 5 ▪ splice site, respectively, and 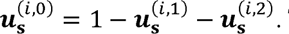 Then,

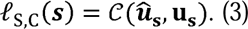

#### Regression loss ℓ_D,R_ in the dual-sequence mode

*ℓ*_D,R_ is used to supervise DeltaSplice’s ability to predict SSU of the target sequence in the dual-sequence mode. Due to the lack of large-scale mutation data during the training process, DeltaSplice randomly samples b pairs of sequences from a batch of data, where b= 48 is the batch size. These sampled sequences simulate the reference sequence, ***s*_r_**, and the target sequence, ***s*_t_**. The SSU of ***s*_r_** and ***s*_t_** in brain tissues are utilized as the reference SSU, denoted as ***u*_*s*_f__**, and the target SSU, denoted as ***u*_*s*_t__**, respectively. Consequently, the regression loss *ℓ*_D,R_(s_t_) is calculated as 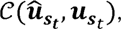 where each bit in 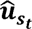 represents the predicted SSU for a site in the target sequence. It is important to note that we opted for random pairs of sequences instead of limiting to orthologous gene pairs, so that the model can contrast gene pairs of varying degrees of similarities.

#### Recovery loss ℓ_D,E_ in the dual-sequence mode

*ℓ*_D,E_ is used to enhance the generalization ability of DeltaSplice. In the dual-sequence mode, when the target sequence is identical to the reference sequence, the predicted target SSU 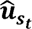 should be the same as the reference SSU ***u*_*s*_r__**. In other words, irrespective of the value of ***u_s_r__***, the model should be capable of recovering the reference SSU when the reference and target sequences are identical. To achieve this, we introduce random reference SSU, which are sampled from a normal distribution. Additionally, *b* pairs of identical sequences are sampled from a batch of data, where *b*=48 is the batch size, to simulate the reference and target sequences. Consequently, *ℓ*_D,E_(s_t_) is denoted as 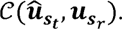

The final loss function used in DeltaSplice can be written as

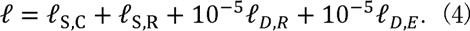

This unified loss function allows for the training of both the single-sequence mode and the dual-sequence mode simultaneously in each training step.

In the training process, the input sequences are shuffled randomly and divided into batches of b=48 sequences for each training step. In each training step, the feature representations for all sequences in the training batch are extracted with the feature-extraction module, and the predictions in the single-sequence mode are computed based on the feature representations. Then, DeltaSplice randomly samples *b* pairs of sequences and their corresponding SSU values. The feature **v_s_t__ - v_s_r__ + v_u_r__** for paired sequences with reference SSU in the dual-sequence mode are computed based on the prior derived feature representations, and thus the predictions in the dual-sequence mode can be calculated. After the computation of all predictions in both the single-sequence mode and the dual-sequence mode, the loss in equation (4) is used to update the model parameters. To update the model parameters, the Adam optimizer is employed. Additionally, a learning rate scheduler is implemented to expedite the convergence process. The initial learning rate is set to 0.002, and the scheduler reduces the learning rate by half after each epoch.

Five models were trained using different random number seeds, and all predictions described in the study used the average of these models.

### Baseline methods

Three state-of-the-art algorithms were used as baseline methods for benchmarking with DeltaSplice in our experiments, including MMSplice (Cheng et al. 2019), SpliceAI (Jaganathan et al. 2019) and Pangolin (Zeng and Li 2022). MMSplice predicts Δlogits of SSU caused by mutations, instead of directly predicting the value of ΔSSU. SpliceAI predicts splice site probabilities, which were used as a proxy of SSU. Pangolin predicts SSU directly. When evaluating the performance of Pangolin in predicting SSU in brain tissues (Figs. 2 and 3), we retrained the Pangolin model using the SSU data estimated from human brain tissue to have a direct comparison with DeltaSplice. For all other evaluations of baseline methods, we used pre-trained models and followed the instructions provided by the original studies.

### Evaluation of lineage-specific splicing changes

For each splice site in the human genes in the test dataset, we identified its orthologous splice site in each of the other seven species. This was done by converting the human coordinate to the coordinates in the other genomes using liftOver (https://genome.ucsc.edu/cgi-bin/hgLiftOver). SSU of the orthologous splice sites was predicted using DeltaSplice and other baseline methods, and the predicted ΔSSU values were compared with experimental measurements from RNA-seq.

### Evaluation of VexSeq data, MFASS data and SNVs in *FAS* exon 6

We obtained Vex-seq (Adamson et al. 2018) dataset from the github repository of MMSplice, which comprises 1,960 mutations and their corresponding PSI values of the reference and alternative alleles, as measured by RNA-seq. For each variant, we used 30,000 bp flanking sequence of the corresponding endogenous gene, and the reference and alternative allele sequences were used for the prediction of ΔPSI by the baseline methods. For DeltaSplice, we also used the PSI of the reference allele as model input. The *FAS* exon 6 dataset (Baeza-Centurion et al. 2019), which consists of 3,072 possible combinations of 12 SNVs and the corresponding PSI values of *FAS* exon 6 as measured by RNA-Seq, was analyzed similarly.

The MFASS (Cheung, et al., 2019) dataset comprises 27,183 SNVs along with their corresponding impacts on exon inclusion. Among these SNVs, 1,048 variants with a Δsplice index::S -0.5 (splice index is conceptually equivalent to PSI, but was derived from a fluorescent reporter readout) were regarded as splicing-disrupting variants (SDVs) in our analysis and classified as positive samples, while the remaining mutations are classified as negative samples. For evaluating the dual-sequence mode of DeltaSplice, we utilized the splice index of the reference allele as an proxy of reference SSU u_r_ for model input.

For DeltaSplice, SpliceAI and Pangolin, the predicted PSI for each cassette exon was calculated as the mean of the predicted SSU of the 3□ or 5□ splice sites of the exon, which was then used to calculate ΔPSI by comparing the alternative and reference alleles. For MMSplice, we followed its code to use the reference PSI to convert the predicted Δlogits into ΔPSI.

### Evaluation of sQTLs in brain tissues

sQTLs from 13 subregions of human brain tissues were obtained from GTEx v8 (Garrido-Martin et al. 2021). Among them, we extracted 7,906 sQTLs within 200 bp of cassette exons and the sQTL exon junctions support the inclusion and skipping isoforms of the cassette exons, respectively. For comparison, we obtained common single nucleotide polymorphisms (SNPs) from the 1000 Genome phase 3 database (The 1000 Genomes Project Consortium 2015) and used a subset of 582,082 SNPs within 200 bp from cassette exons as background. For sQTLs and background SNPs mapped to multiple cassette exons, we kept only one representative variant-exon pair for each variant selected using the following criteria:

- If the variant overlaps exon A, while it is in the intronic regions of exon B, we kept exon A.
- If the variant was annotated as an intronic variant in multiple variant-to-exon pairs, we chose the one with minimal distance between the variant and the exon.
- If there are still more than one variant-to-exon pairs with the same minimal distance, we select the exon with maximal evidence based on mRNA/EST sequences.

When predicting the impact of sQTLs with DeltaSplice, the dual-sequence mode is used with the RNA-seq derived SSU as a reference. Note that common SNPs might include sQTLs, so the enrichment of splicing-altering variants among sQTLs reported in this study might represent an underestimation.

### Evaluation of *de novo* mutations related to neurodevelopmental disorders

We collected a comprehensive list of *de novo* mutations (DNMs) of neurodevelopmental disorders (NDD) patients and corresponding controls (e.g., unaffected siblings when available), which became available prior to 2020 from different data sources, including denovo-db database (v.1.6.1) (Turner et al. 2017), Simons Simplex Collection (An et al. 2018), and literature survey including a large-scale exome sequencing study of autism (Satterstrom et al. 2020). Mutations obtained by the whole genome sequencing (WGS) dataset and whole exome sequencing (WES) dataset were analyzed separately. Studies that only reported DNMs for a subset of genes (i.e. by targeted sequencing analysis) were excluded from our analysis. To remove redundancy from different studies, we merged mutations from samples that share more than 50% of their DNMs (and have the same phenotypes if phenotype information is available) into a single sample. After redundancy removal, we obtained a total of 3,564 cases and 2,162 controls for the WGS dataset, and 11,280 cases and 1,880 controls for WES dataset (Supplemental Tables S4 and S5). Note that we used the combination of sample ID and variant ID to identify each DNM for enrichment analysis, to allow recurrent mutation events in the same nucleotide position, although such events are expected to be very rare.

We then associated each DNM with human exons <5000 bp from the DNM. For variants that can be associated with multiple exons, one variant-to-exon pair was selected to predict the impact of the DNM on splicing using similar criteria as described above.

To identify splicing-altering mutations, we excluded DNMs that disrupt splice sites (GU/AG di-nucleotides) or lead to stop gain/loss, since these likely gene-disrupting (LGD) mutations were previously associated with autism. For the dual-sequence mode of DeltaSplice, the RNA-seq derived SSU was used as a reference.

### Code Availability

DeltaSplice source code and documentation are available at https://github.com/chaolinzhanglab/DeltaSplice.

## SUPPLEMENTAL FIGURE LEGENDS

**Figure S1: Distribution of human SSU predicted by DeltaSplice and baseline methods in comparison with experimental measurements by RNA-seq.** Related to Fig. 2A in the main text, but the distribution of predicted SSU values is shown using violin plots for splice sites with different ranges of SSU.

**Figure S2: Distribution of predicted and experimental ΔSSU for splice sites between the other species.** Related to Fig. 3A in the main text. Distribution of ΔSSU predicted by different methods and experimental ΔSSU is shown for splice sites in additional species homologous to humans.

**Figure S3: The performance of DeltaSplice and baseline methods in predicting splicing-altering mutations as measured by reporter assays.** Similar to Fig. 4A in the main text, but for the *FAS* exon 6 dataset.

**Figure S4: The performance of DeltaSplice and baseline methods in predicting splicing-altering mutations at sQTLs.** Similar to Fig. 5 in the main text, but variants overlapping with splice sites were excluded for analysis.

## Notes

### Competing Interest Statement

The authors have declared no competing interest.

### Summary of Updates

Code availability is provided in the revised version.

