## Supplementary figures and images for "Reference-informed prediction of alternative splicing and splicing-altering mutations from sequences"

### Supplemental Figures S1-S4

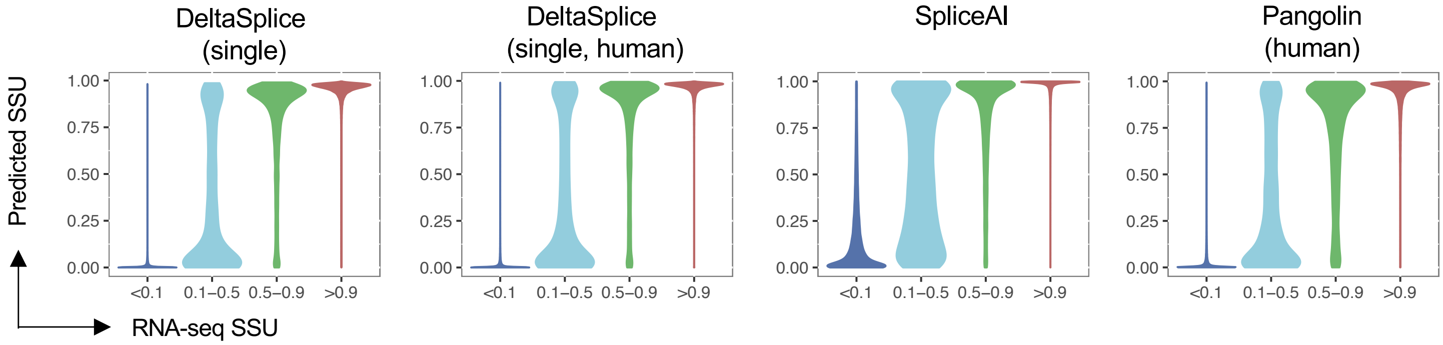

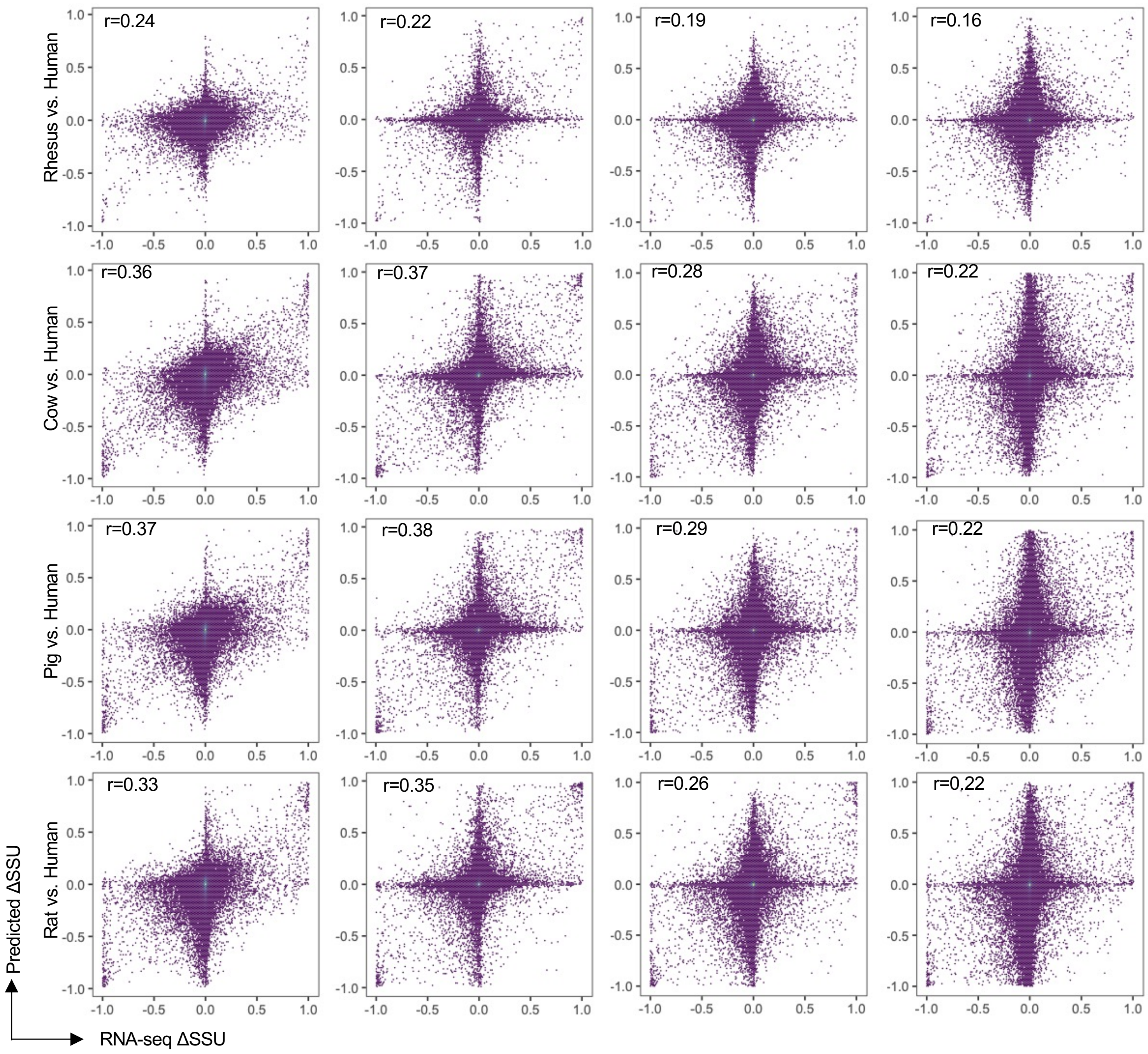

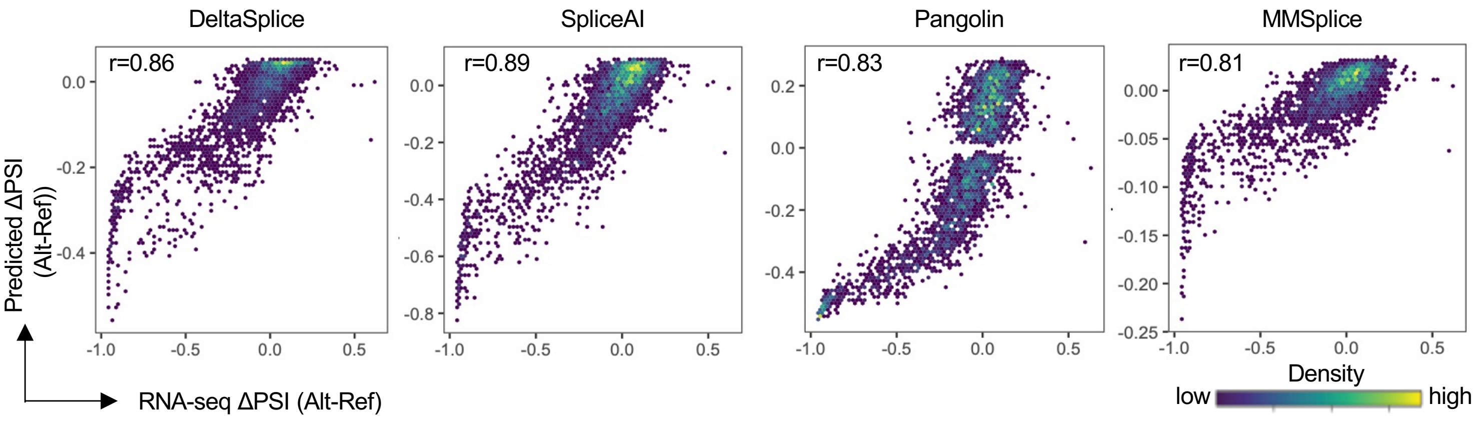

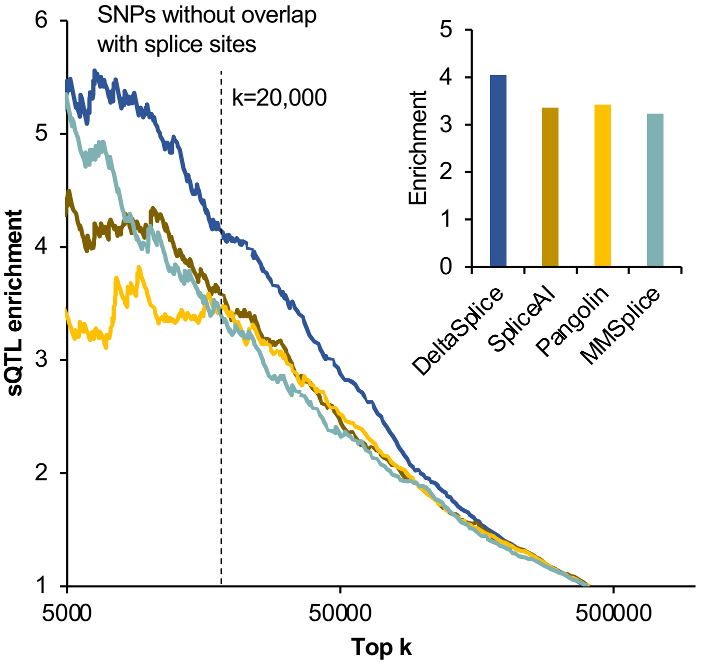
